# Long-term impacts on estuarine benthic assemblages 4.2 years after a mine tailing spill in SE Brazil

**DOI:** 10.1101/2023.08.25.554820

**Authors:** Gabriel C. Coppo, Fabrício A. Gabriel, Ana Carolina A. Mazzuco, Hermano M. Queiroz, Diego Barcellos, Tiago O. Ferreira, Angelo F. Bernardino

## Abstract

The Rio Doce estuary was critically impacted in 2015 by the world’s largest mining tailing spill, with still unclear long-term effects. Here we present a long-term (2015 to 2020) assessment of estuarine benthic assemblages, where Annelida and Mollusca were dominant (52.6% and 26.3%, respectively). Our results demonstrated that the density and richness of benthic taxa decreased in response to an increase in potentially toxic elements concentrations, suggesting a chronic pollution effect that lasted for at least 4.2 years in the estuary. Our data demonstrated that despite the decline in sediment potentially toxic elements concentration over time, there was a continued low habitat quality for the benthic fauna characterized by a reduction of 96% on macroinvertebrate density and 48% on species richness. The long-term impacts on benthic macrofauna highlight that water quality levels can misguide impact assessment programs, and reveal that mine tailings impacts persist for many years in estuarine ecosystems.

## Introduction

In November 2015, 43 million m^3^ of mine tailings were spilled after the collapse of Fundão dam causing massive destruction in SE Brazil (Do Carmo et al., 2017). Over the following 16 days, the tailings were transported for over 600 km along the Rio Doce river basin and reached the estuary and the Atlantic Ocean (Gomes et al., 2017). The event was recognized as the largest environmental disaster in Brazilian history and one of the largest in the world involving mining activity (Almeida et al., 2018; Aires et al., 2018). The volume of the tailings caused an immediate increase in water turbidity and significantly affected riverine sediments, riparian soils and vegetation, and impacted the aquatic fauna of the Rio Doce estuary (Gomes et al., 2017; Bernardino et al., 2019; Queiroz et al., 2018). Once in the estuary, the mine tailings were enriched with potentially toxic elements (PTE) when compared to samples from pre-impact conditions, resulting in significant changes in the soil geochemistry and losses of the local biota (Gomes et al., 2017; Queiroz et al., 2018; Bernardino et al., 2019; Andrades et al., 2020; Sá et al., 2021; Silva et al., 2023). The damage to the estuarine ecosystems was significant, including contamination and sublethal effects on the aquatic fauna, offering ecological risks and deleterious effects to benthic organisms (Gabriel et al., 2020a; 2020b; Weber et al., 2020, Queiroz et al., 2021a).

Multiple lines of evidence demonstrated that the release of mine tailings in the Rio Doce basin were the cause of a significant increase in the soil and water content of potentially toxic elements (PTE) in the estuary; which were not present at similar levels during pre-impact assessments (Queiroz et al., 2018; Queiroz et al., 2021a; 2021b; Queiroz et al., 2022). As a result, the contamination of estuarine soils by bioavailable fractions of PTE was the main threat to the Rio Doce aquatic ecosystems (Gabriel et al., 2020a; 2020b; Queiroz et al., 2021a; 2021b; Queiroz et al., 2022). Potentially toxic elements contamination is particularly relevant and a problem of increasing significance for ecological, evolutionary, nutritional, and environmental reasons (Jaishankar et al., 2014). Marine and benthic organisms show different tolerant capacities to PTE accumulation, but once they are incorporated in aquatic food webs, there are increasing risk to human health through fish and seafood ingestion (Gnandi et al., 2006; Lahijanzadeh et al., 2019). Trace element toxicity is recognized as one of the main threats to aquatic ecosystems worldwide (Zhou et al., 2008) and, therefore, require their long-term monitoring due to their mobility and toxicity risks (Kraus and Wiegand 2006).

The disaster’s enormous scale meant, however, that large soil and sediment areas remain residually contaminated and that patches of elevated contamination may have remained in the Rio Doce estuary (Gabriel et al., 2020a; Queiroz et al., 2021a; 2021b). This is significant because many studies and monitoring programs have found that water and aquatic sediment properties in many localities of the Rio Doce basin were considered “safe”, and therefore the disaster’s footprint would have ceased (Richard et al., 2020; Abessa et al., 2023). Although water and sediment quality parameters in upper river streams may have returned to pre-impact conditions, estuarine soils and marine sediments may act as long-term sources of potentially toxic elements due to local soil redox conditions (Queiroz et al., 2018). The strong sink effect has been widely demonstrated in the Rio Doce estuary where significant (meaning above pre-impact levels) concentrations of metals and metalloids remained in soils for many years (Gabriel et al., 2021), with significant environmental risks and chronic pollution effects on the benthic fauna (Vermeulen, 1995; Riba et al., 2002; Gabriel et al., 2020a; Queiroz et al., 2022; Coppo et al., 2023). Understanding the long-term pollution effects pollution on benthic assemblages are therefore of great importance for contamination assessments in the Rio Doce estuary as these assemblages are excellent bioindicators of estuarine health (Muniz et al., 2002; Venturini et al., 2004; Barcellos et al., 2021).

Estuarine sediments contains a diverse and sensitive benthic fauna, which play crucial ecological functions, such as in sedimentary irrigation, nutrient cycling in the sediment-water interface, organic matter decomposition, and energy transfer to trophic levels higher (Snelgrove et al., 2000; Kristensen et al., 2012; Kristensen et al., 2014). The benthic macrofauna lives in close association with sediments and have short generation periods and sedentary characteristics; characters that make them useful to impact assessments in aquatic habitats (Warwick and Clarke 1993; Hyland et al., 2005). In addition, benthic species have different tolerance levels to contaminants and sedimentary quality conditions, which are useful to understand anthropogenic impacts over multiple spatial and temporal scales (Muniz et al., 2011; Hadlich et al., 2018). Therefore, we can expect that chemical disturbances will greatly induce changes in benthic community assemblages with subsequent impacts on ecosystem dynamics and functions for long periods of time (Clements et al., 2000; Burd, 2002; Lancellotti and Stotz, 2004).

In this context, our study evaluated the sedimentary contamination and benthic organisms of the Rio Doce estuary with the aim to provide the first assessment of long-term trends of contaminants and assemblage succession in the area. Although a number of studies have provided information of chemical sedimentary concentrations and behavior of potentially toxic elements in soil and water (Gabriel et al., 2020b; Sá et al., 2021), there is relatively limited information of long-term impacts of tailings in benthic assemblages (Burd, 2002; Sweetman et al., 2020). Current progress in the study of the Rio Doce mine tailings impact (2-3 yrs after impact) have showed that following the initial deposition we could observe a chronic effect in benthic meiofaunal (<300 µm) and demersal fish estuarine assemblages due to PTE contamination and resuspension (Bernardino et al., 2019; Andrades et al., 2021; Coppo et al., 2023). Given that there is a potential for long-term (>3 years) contamination of estuarine soils when compared to baseline conditions (Gabriel et al., 2021), here we evaluated the benthic macrofaunal assemblage in the Rio Doce estuary before the disaster and up to 4.2 years after the arrival of the mining tailings. By comparing the macrofaunal succession after the initial tailing deposition, with temporal soil contamination, we hypothesized that a decrease in potentially toxic element concentrations would be followed by a recolonization and gradual recovery of invertebrate assemblages that were observed before the impacts.

## Material and Methods

### Study area and sampling strategy

The Rio Doce estuary is located on the southeast coast of Brazil, in the Eastern Marine Ecoregion categorized as a humid tropical area, with a rainy season from October to March and a dry season from April to September (Figure 1; Bernardino et al., 2015). The Rio Doce estuary is characterized by a highly dynamic main channel, with river discharge being the main cause of water circulation, causing low salinity levels (Gomes et al., 2017; Sá et al., 2021).

**Figure 1.**
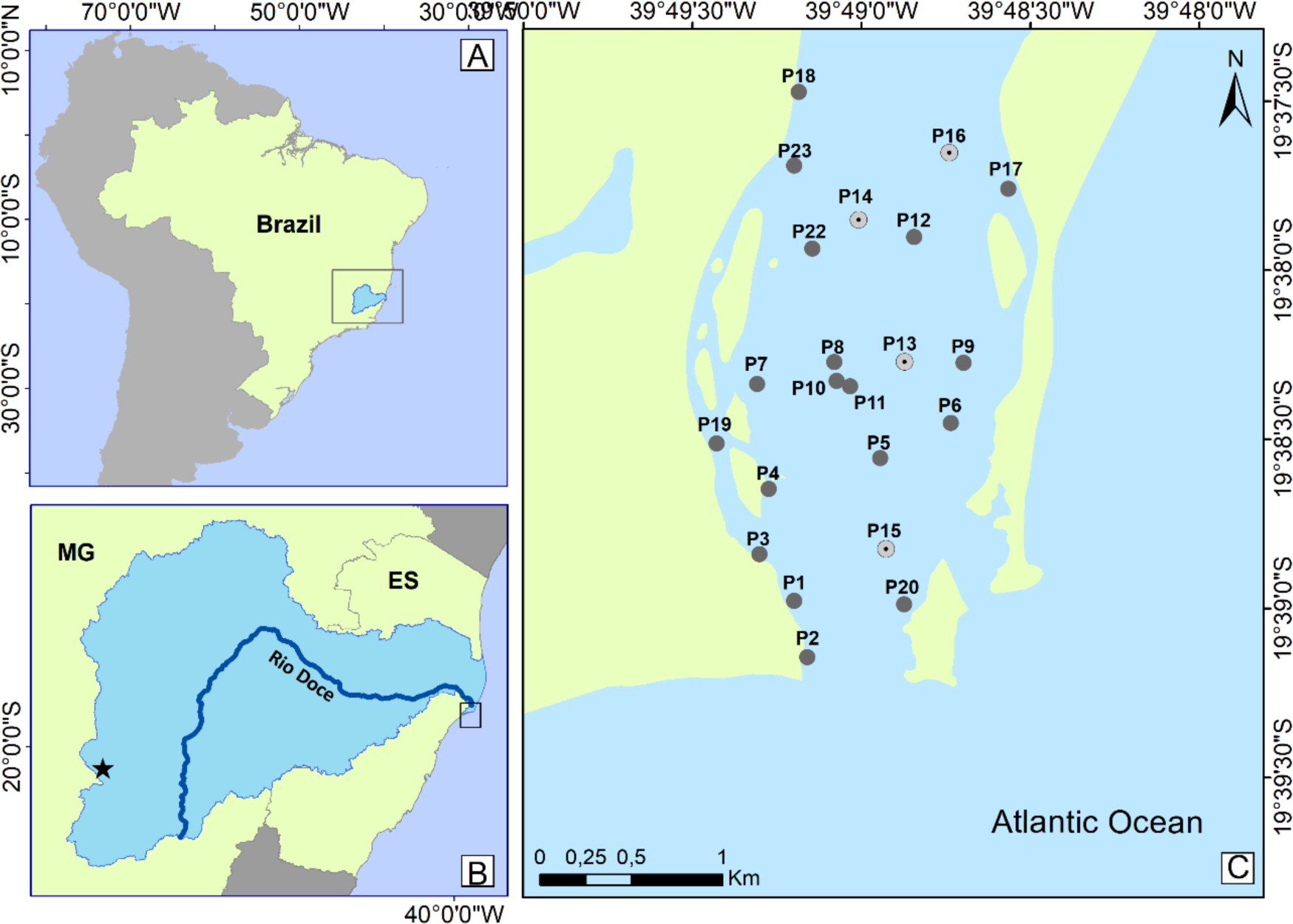
Map of the Rio Doce estuary indicating the 17 random sampling stations monitored during this study. The light gray stations (#13, 14, 15 and 16) were sampled by Gomes et al. (2017) and used to determine pre-impact sediment contamination levels.

Sampling of benthos (>500µm) and sediments in the estuary followed a randomized BACI (Before-After Community Impact; Underwood, 1992) approach along 17 stations in the estuary (Figure 1). The initial impacts on the benthic macrofauna in the Rio Doce estuary (pre- and after-impacts, scale of days) were demonstrated by Gomes et al., (2017). By using identical sampling and analytical protocols, we incorporated the data of Gomes et al., (2017) and expanded the biological and sedimentary sampling in the estuary up to 4.2 years after the disaster (August 2017, January 2018, August 2018, February 2019, and January 2020; Figure 2). Sediment sampling was carried using a Van Veen Grab sampler with three replicates per station. In each replicate, the top 5 cm of undisturbed surface sediments were sub-sampled for grain size, potentially toxic elements concentrations, and another acrylic tube of 10 cm (internal diameter) was used to sample the benthic macrofauna. The macrobenthos samples were fixed in 10% formaldehyde after sampling without sieving. The sediment samples for potentially toxic element determination were stored in sterile containers (HNO_3_ 10% v/v) and immediately preserved on ice and later frozen at −20 °C. Environmental conditions (water temperature, pH, salinity, and total dissolved solids-TDS) were measured at the time of sampling using a portable HANNA multi-parameter (HI9829).

**Figure 2.**
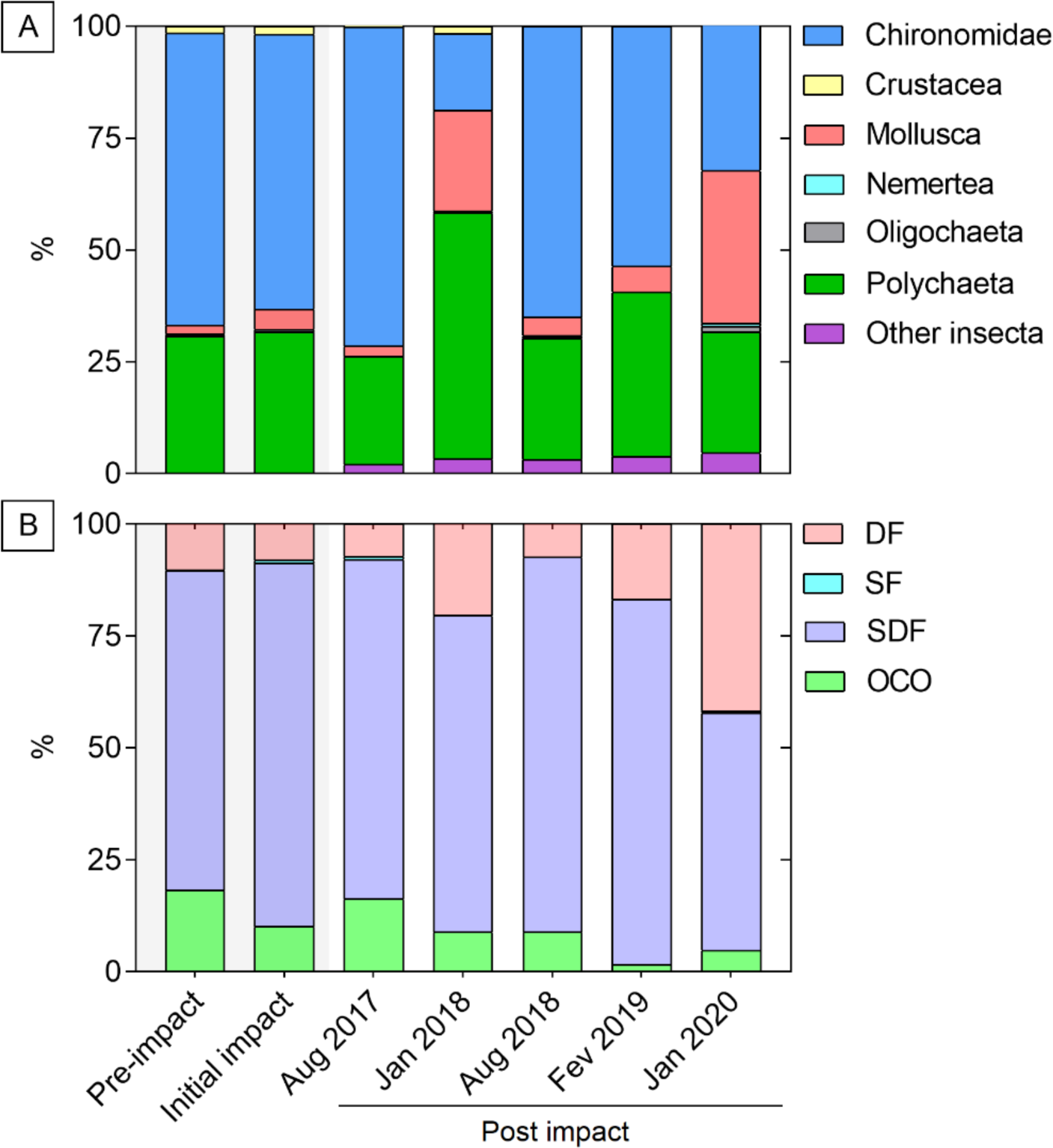
Relative abundance of macrofauna assemblages in the Rio Doce estuary: (A) taxonomic composition, and (B) trophic groups.

### Laboratory procedures

The sediment samples were analyzed for the PTE content using the tri-acid method according to the USEPA 3052 protocol (USEPA, 1997). Sediment samples were digested (100 g of wet weight) in a microwave with 2 mL of HNO_3_ (65%), 6 mL of HCl (37%), and 2 mL of HF (40%) for 40 min. After digestion, the extracts were filtered (0.8 µm Millipore membrane) according to the USEPA 6010C protocol, to determine multiple analyte elements using inductively coupled plasma optical emission spectroscopy (ICP-OES, Thermo Fisher Scientific, Waltham, MA, USA). The EPA 3051A method validation for PTE analysis was performed by comparing the results to the values of certified reference material NIST SRM 2709a.

The sediments grain size distribution was performed using the pipette method with previous treatment for oxidation of the organic matter with H_2_O_2_ and dispersion using a combination of physical (overnight shaking) and chemical [0.015 mol L^-1^ (NaPO_3_)_6_ + 1.0 mol L^-1^ NaOH] (Gee and Bauder, 1986). The organic matter content was determined by loss-on-ignition (Dean, 1974), which consists of adding approximately 10 g of sediment sample in crucible and then heating for 4 h in a muffle furnace at 550 °C. The benthic macrofaunal samples were washed and sieved in a 500 µm mesh sieve, and the retained material transferred and preserved in 70% ethanol for further sorting and identification. All macro-invertebrates were counted and identified under a stereomicroscope up to Family or lower taxonomic levels using taxonomic keys (e. g., Amaral and Nonato, 1996; Amaral et al. 2005). Functional groups (suspension feeder - SF; detritivore feeder - DF; surface deposit feeder - SDF; omnivores, carnivores and other - OCO) were determined and grouped according to Jumars et al. (2015).

### Statistical analyses

The benthic macrofaunal data was grouped in two treatments - pre-impact (November 2015; N=10 replicates), initial impact (0 to 2 days after the tailings arrival; 11 replicates), and post-impact (17 replicates per sampling) up to 4.2 years after the spill (August 2017, January 2018, August 2018, February 2019, and January 2020). For each group of macrofaunal samples, we calculated the mean assemblage ecological metrics (density, abundance, richness, Shannon diversity index, and Pielou evenness index) (Margalef, 1958; Fisher et al., 1943; Hurlbert, 1971). In addition, we estimated the macrofaunal assemblage functioning by their indices of functional richness (FRic), functional dispersion (FDis), functional evenness (FEve), and entropy (FRaoQ; Mason et al., 2005). FRic indicates the amount of niche space filled for the character in a community, based on the niche space filled by the species within the community and the absolute range of the character. The FEve index may be interpreted as the degree to which the abundance of a community is distributed in niche space, where lower values of FEve indicate that the niche space is under-utilized (Mason et al., 2005). The indices FDis and RaoQ are used to quantify how functionally similar are the individuals spatially (Botta-Dukát, 2005).

The sedimentary concentration of potentially toxic elements, granulometry, and water physical-chemical variables were compared among three different impact conditions (pre-impact, initial impact, and post impact) by an Permutational Analysis of Variance (PERMANOVA; Anderson, 2017). The PERMANOVA was also performed to compare biological metrics (density, richness, diversity, evenness, FRic, FDis, FEve, and FRao) among sampling events. Permutational Multivariate Analysis of Variance (PERMANOVA) was applied for multiple variables comparisons among impact conditions (macrofaunal assemblage composition). Additionally, we used PERMANOVA to tested spatial differences in macrofauna assemblage between sampling stations for the post-impact campaigns. Data was transformed when needed to fit these premises (log(x+1), log(x+10)). A canonical analysis of principal coordinate (CAP; Anderson and Willis, 2003) was performed to determine the association between PTE contamination, sediment grain size, and benthic assemblage across different impact levels (pre-impact, initial impact, and post impact events). PERMANOVA and CAP were based on a Bray-Curtis distance resemblance matrix under a reduced residuals model and data was firstly transformed (square-root) (Clarke and Gorley, 2006). Samples with total density equal to zero were excluded when required by the analysis (assemblage composition, FRic, RaoQ, and CAPs). All graphic design and analysis were performed using R Environment (R Core Team, 2021), and significant differences were considered when p < 0.05.

## Results

### Environmental conditions

The sediment grain size remained similar over time, with high contents of sand (from 79.7 ± 20.9% in August 2017 to 95.6 ± 3.0% in January 2020; Table 1) in most sites and a lower fine fractions content (silt and clay varying from 4.4 ± 3.0% in January 2020 to 20.3 ± 20.9 % in August 2017; Table 1) (F = 1.97, p = 0.11; Sand, F = 1.95, p = 0.08; Table 1). Total sedimentary organic matter varied from 5.3% ± 4.8% in August 2017 to 4.0% ± 3.2% in January 2018, without significant differences between the periods (F = 0.32, p = 0.86; Table 1). Water salinity at time of sampling ranged from 0.7 ± 0.9 in August 2017 to 0.1 ± 0.1 in February 2019. The sampling in August 2017 presented the highest values of total dissolved solids (561 ± 826.6 mg/L), decreasing over time and reaching an average concentration of 53.1 ± 44.2 mg/L in January 2020 (Table 1). The total sediment organic matter and TDS data were not compared to pre-impact conditions due to the absence of data.

**Table 1.** Summary of water column (pH, temperature, salinity, and total dissolved solids - TDS) and sediment (silt+clay, sand, organic matter - OM, and potentially toxic elements - PTE concentrations) conditions to the Rio Doce estuary in pre-impact, initial impact, and post impact. Values represent average ± SD, granulometry and OM in %, and metal(loid)s concentrations in mg kg^-1^. *Pre-impact and initial impacts assessments were defined in Gomes et al. (2017).

|  |  | Pre-impact* | Initial impact* | Post impact |  |  |  |  |
| --- | --- | --- | --- | --- | --- | --- | --- | --- |
|  |  |  |  | Aug 2017 | Jan 2018 | Aug 2018 | Fev 2019 | Jan 2020 |
| Water | Salinity | 7.0 ± 2.6 | 1.3 ± 1.6 | 0.7 ± 0.9 | 0.1 ± 0.1 | 0.1 ± 0.2 | 0.1 ± 0.1 | 0.1 ± 0.04 |
|  | Temp.<br>(°C) | 28.6 ± 0.7 | 27.7 ± 0.5 | 23.1 ± 0.7 | 30.3 ± 1.3 | 25.3 ± 0.9 | 30.5 ± 0.9 | 29.1 ± 0.2 |
|  | pH | 8.0 ± 0.3 | 7.8 ± 0.4 | 7.4 ± 0.4 | 7.1 ± 0.4 | 7.8 ± 0.4 | 8.7 ± 0.8 | 9.1 ± 0.2 |
|  | TDS | - | - | 561.0 ± 826.6 | 95.29 ± 74.7 | 109.4 ± 80.8 | 139.8 ± 73.9 | 53.1 ± 44.2 |
| Sediment | Silt+Clay | 18.9 ± 25.3 | 7.8 ± 7.4 | 20.3 ± 20.9 | 13.6 ± 11.5 | 16.6 ± 12.2 | 11.3 ± 23.3 | 4.4 ± 3.0 |
|  | Sand | 81.1 ± 25.3 | 92.2 ± 7.2 | 79.7 ± 20.9 | 86.4 ± 11.5 | 83.4 ± 12.2 | 88.7 ± 23.3 | 95.6 ± 3.0 |
|  | OM | - | - | 5.3 ± 4.7 | 4.0 ± 3.2 | 4.7 ± 3.2 | 4.7 ± 3.3 | 4.3 ± 2.9 |
|  | As | 3.3 ± 0.9 | 3.8 ± 2.6 | 7.8 ± 8.1 | 1.8 ± 1.9 | 5.1 ± 2.9 | 5.0 ± 3.6 | 4.0 ± 2.4 |
|  | Cd | 0 | 0 | 3.6 ± 1.5 | 1.1 ± 0.6 | 1.6 ± 0.6 | 0.0 | 0.1 ± 0.1 |
|  | Co | 0.3 ± 0.3 | 0 | 10.0 ± 3.4 | 3.3 ± 1.1 | 7.0 ± 1.4 | 5.4 ± 6.2 | 4.0 ± 5.2 |
|  | Cr | 3.6 ± 2.3 | 7.5 ± 8.1 | 47.4 ± 15.7 | 9.8 ± 7.0 | 25.9 ± 9.3 | 11.1 ± 12.8 | 14.0 ± 15.8 |
|  | Cu | 1.3 ± 0.5 | 2.2 ± 2.5 | 9.4 ± 4.1 | 2.7 ± 1.9 | 3.6 ± 1.8 | 5.0 ± 6.4 | 4.9 ± 7.0 |
|  | Fe | 9025.8 ± 2923.7 | 15201.1 ± 12098.0 | 34333.3 ± 10162.3 | 9252.8 ± 3598.8 | 16372.5 ± 4786.9 | 9900.9 ± 10094.3 | 8692.9 ± 9485.2 |
|  | Mn | 230.9 ± 92.5 | 274.2 ± 132.3 | 539.8 ± 227.3 | 181.0 ± 63.6 | 308.4 ± 100.9 | 167.9 ± 123.9 | 142.9 ± 102.7 |
|  | Ni | 2.1 ± 0.5 | 3.1 ± 2.7 | 15.4 ± 5.2 | 2.1 ± 1.1 | 9.8 ± 2.3 | 5.6 ± 6.1 | 5.9 ± 7.9 |
|  | Pb | 4.7 ± 0.9 | 4.0 ± 0.5 | 100.2 ± 48.9 | 2.9 ± 1.1 | 6.5 ± 1.5 | 95.5 ± 61.3 | 135.4 ± 73.9 |
|  | Zn | 1.6 ± 0.2 | 2.1 ± 1.7 | 38.7 ± 14.9 | 11.0 ± 6.0 | 25.5 ± 9.0 | 13.7 ± 13.4 | 12.5 ± 14.2 |

The concentrations of potentially toxic elements in sediments exhibited a significant increase from the pre-impact to the post impact periods (Table 1). Based on the baseline values from the pre-impact (As = 3.3 ± 0.9 mg kg^-1^; Cd = 0 mg kg^-1^; Co = 0.3 ± 0.3 mg kg^-1^; Cr = 3.6 ± 2.3 mg kg^-1^; Cu = 1.3 ± 0.5 mg kg^-1^; Fe = 9025.8 ± 2923.7 mg kg^-1^; Mn = 230.9 ± 92.5 mg kg^-1^; Ni = 2.1 ± 0.5 mg kg^-1^; Pb = 4.7 ± 0.9 mg kg^-1^; Zn = 1.6 ± 0.2 mg kg^-1^; Table 1), the higher increases in PTE were observed in August 2017 (21 months after the disaster) for Cd (mean of 3.6 mg kg^-1^; 360-fold), Zn (mean of 38.7 mg kg^-1^; 24-fold), and Co (mean of 10 mg kg^-1^; 20-fold) (Cd; F = 60.9, p < 0,0001, Zn; F = 19.7, p < 0.001, Co; F = 10.8, p = 0.01; Table 2); while Pb reached the highest level in January 2020 (mean of 135.4 ± 73.9 mg kg^-1^; 29-fold), which is significantly greater when compared to the pre-impact conditions (Feb 2019; F = 63.1, p = 0.023, Jan 2020; F = 63.1, p < 0.001). Cd and Mn concentrations gradually decreased until January 2020, reaching values slightly below pre-impact conditions.

**Table 2.** Mean density (ind·m^−2^) and relative abundances of macrofaunal assemblages (Chironomid larvae removed) in each assessment (pre-impact, initial impact, and post impact) in the Rio Doce estuary. P = Polychaeta, M = Mollusca, C = Cumacea, I = Insecta; Feeding types: SDF = Surface deposit feeder, DF = Detritivore feeder, OCO = Omnivores, carnivore and other feeder, SF = Suspension feeder. *Pre-impact and initial impacts assessments were defined in Gomes et al. (2017).

| Species | Pre-impact* |  | Initial impact* |  | Post impact |  |  |  |  |  |  |  |  |  |
| --- | --- | --- | --- | --- | --- | --- | --- | --- | --- | --- | --- | --- | --- | --- |
|  |  |  |  |  | August 2017 |  | January 2018 |  | August 2018 |  | February 2019 |  | January 2020 |  |
|  | Mean m<sup>2</sup> | Rel % | Mean m<sup>2</sup> | Rel % | Mean m<sup>2</sup> | Rel % | Mean m<sup>2</sup> | Rel % | Mean m<sup>2</sup> | Rel % | Mean m<sup>2</sup> | Rel % | Mean m<sup>2</sup> | Rel % |
| **Annelida** |  |  |  |  |  |  |  |  |  |  |  |  |  |  |
| *Amphicteis* sp. (P, SDF) | 391.7 (440) | 11.9 | 450 (595) | 11.5 | 0 | 0 | 0 | 0 | 0 | 0 | 0 | 0 | 0 | 0 |
| *Capitella capitata* (P, SDF) | 50 (90) | 1.5 | 79.2 (127) | 2.0 | 0 | 0 | 0 | 0 | 0 | 0 | 0 | 0 | 4.6 (8) | 3.3 |
| *Cirrophorus* sp. (P, SDF) | 33.3 (58) | 1.0 | 33.3 (60) | 0.9 | 0 | 0 | 1.5 (3) | 1.6 | 0 | 0 | 0 | 0 | 0 | 0 |
| *Glycera* sp. (P, DF) | 204.2 (261) | 6.2 | 0 | 0 | 0 | 0 | 0 | 0 | 6.8 (12) | 3.8 | 4.9 (9) | 5.6 | 0 | 0 |
| *Heteromastus similis* (P, SDF) | 37.5 (68) | 1.1 | 0 | 0 | 0 | 0 | 0 | 0 | 0 | 0 | 0 | 0 | 0 | 0 |
| *Isolda pulchella* (P, SDF) | 425 (540) | 12.9 | 929.2 (1220) | 23.7 | 0 | 0 | 0 | 0 | 0 | 0 | 0 | 0 | 0 | 0 |
| *Laeonereis* sp. (P, SDF) | 1050 (905) | 31.8 | 1325 (1510) | 33.8 | 169.1 (216) | 73.6 | 44.8 (62) | 48.2 | 118.9 (176) | 66.5 | 67.7 (79) | 78.1 | 48 (50) | 34.3 |
| *Magelona* sp (P, SDF) | 91.7 (153) | 2.8 | 0 | 0 | 2 (4) | 0.9 | 0.5 (1) | 0.5 | 6.3 (12) | 3.5 | 0 | 0 | 0 | 0 |
| *Mediomastus* sp. (P, DF) | 16.7 (30) | 0.5 | 8.3 (16) | 0.2 | 0 | 0 | 3.9 (8) | 4.2 | 0 | 0 | 0 | 0 | 0 | 0 |
| *Allita succinea* (P, OCO) | 525 (543) | 15.9 | 395.8 (425) | 10.1 | 37.5 (56) | 16.3 | 8.3 (13) | 8.9 | 14.2 (24) | 7.9 | 1.5 (3) | 1.7 | 6.7 (12) | 4.8 |
| *Pholoe* sp. (P, OCO) | 25 (47) | 0.8 | 0 | 0 | 0 | 0 | 0 | 0 | 0 | 0 | 0 | 0 | 0 | 0 |
| *Polydora* sp. (P, SDF) | 25 (45) | 0.8 | 0 | 0 | 0 | 0 | 5.4 (10) | 5.8 | 4.2 (8) | 2.4 | 1 (2) | 1.1 | 0.5 (1) | 0.4 |
| *Sigambra grubii* (P, OCO) | 29.2 (54) | 0.9 | 0 | 0 | 0 | 0 | 0 | 0 | 1.6 (3) | 0.9 | 0 | 0 | 0 | 0 |
| *Sthenelais* sp. (P, OCO) | 20.8 (38) | 0.6 | 0 | 0 | 0 | 0 | 0 | 0 | 0 | 0 | 0 | 0 | 0 | 0 |
| Oligochaeta (O, SDF) | 41.7 (69) | 1.3 | 45.8 (83) | 1.2 | 0 | 0 | 0.5 (1) | 0.5 | 1.1 (2) | 0.6 | 0 | 0 | 2.6 (5) | 1.8 |
| **Mollusca** |  |  |  |  |  |  |  |  |  |  |  |  |  |  |
| *Tellina* sp. (M, SDF) | 50 (87) | 1.5 | 62.5 (108) | 1.6 | 0 | 0 | 1 (2) | 1.0 | 0 | 0 | 0 | 0 | 2.1 (4) | 1.5 |
| *Tagelus plebeius* (M, SF) | 4.2 (8) | 0.1 | 25 (48) | 0.6 | 1.6 (3) | 0.7 | 0 | 0 | 0 | 0 | 0 | 0 | 0.5 (1) | 0.4 |
| *Macoma* sp. (M, SD) | 8.3 (16) | 0.3 | 62.5 (104) | 1.6 | 1.6 (3) | 0.7 | 10.2 (19) | 11.0 | 16.8 (32) | 9.4 | 1.9 (4) | 2.2 | 15.5 (19) | 11.1 |
| *Melanoides tuberculata* (M, SDF) | 8.3 (16) | 0.3 | 0 | 0 | 0.4 (1) | 0.2 | 0 | 0 | 0 | 0 | 0 | 0 | 0 | 0 |
| *Heleobia australis* (M, DF) | 116.7 (187) | 3.5 | 308.3 (426) | 7.9 | 16.7 (26) | 7.3 | 15.1 (23) | 16.2 | 6.3 (12) | 3.5 | 9.7 (15) | 11.2 | 57.8 (64) | 41.3 |
| **Arthropoda** |  |  |  |  |  |  |  |  |  |  |  |  |  |  |
| *Diastylis* sp. (C, SDF) | 137.5 (213) | 4.2 | 187.5 (279) | 4.8 | 0.8 (2) | 0.3 | 1.9 (4) | 2.1 | 0 | 0 | 0 | 0 | 0 | 0 |
| Chironomidae (I) | 6216.7 (8105) |  | 6229.2 (6664) |  | 616.4 (753) |  | 20 (28) |  | 364.7 (366) |  | 109.6 (120) |  | 71.2 (68) |  |
| Other insects | 0 |  | 0 |  | 18.3 (29) |  | 3.9 (7) |  | 17.4 (26) |  | 7.8 (12) |  | 10.3 (15) |  |
| **Nemertea** | 8.3 (16) | 0.3 | 8.3 | 0.2 | 0 | 0 | 0 | 0 | 2.6 (5) | 1.5 | 0 | 0 | 1.5 (3) | 1.1 |
| **Total individuals** | 2284 |  | 2436 |  | 2168 |  | 240 |  | 1066 |  | 419 |  | 429 |  |
| Richness (S) | 23 |  | 15 |  | 10 |  | 13 |  | 12 |  | 8 |  | 12 |  |
| Shannon (H') | 1.10 (0.4) |  | 0.89 (0.4) |  | 0.51 (0.4) |  | 0.36 (0.4) |  | 0.48 (0.5) |  | 0.50 (0.4) |  | 0.94 (0.4) |  |
| Evenness (Pielou 1969) | 0.71 (0.2) |  | 0.62 (0.2) |  | 0.72 (0.2) |  | 0.79 (0.2) |  | 0.70 (0.3) |  | 0.78 (0.2) |  | 0.84 (0.1) |  |
| Dominance | 0.45 (0.2) |  | 0.54 (0.2) |  | 0.70 (0.3) |  | 0.78 (0.3) |  | 0.72 (0.3) |  | 0.70 (0.3) |  | 0.47 (0.2) |  |

### Macrofaunal assemblages

The total macrofauna density showed a 96% decrease in the post impact condition when compared to the pre-impact and after the initial impact (F = 62.5, p < 0.001; Table S1). The density of chironomid larvae did not show differences over the post impact condition (F = 2.0, p = 0.095), but was on average 96% lower when compared to pre-impact and initial impact (F = 10.8, p < 0.001; Table S3). During the impact assessments, 4,322 individuals from 19 taxa were identified at the 17 sampling sites in the Rio Doce estuary (Table 2). We observed significant changes in benthic macrofaunal assemblages in each of the sampling campaigns (PERMANOVA F=40.3; p<0.001; Table S2). Annelida and Mollusca were the most frequent taxa, corresponding respectively to 52.6% and 26.3% of all benthic macroinvertebrates in this study (Figure 2). We observed a similar relative abundance of major feeding guilds of benthic invertebrates in the estuary across all periods sampled, with occasional higher proportion of mollusks during the wet seasons (January 2018 and 2020). Chironomids were highly abundant in all times sampled, along with *Laeonereis* sp. (Polychaeta), *Heleobia australis* (Mollusca), and *Alitta succinea* (Polychaeta). It is noteworthy that *Heteromascus similis*, *Pholoe* sp., and *Sthenelais* sp. were not detected after the initial impact; *Amphicteis* sp., and *Isolda pulchella* were not detected post impact; and *Glycera* sp., *Magelona* sp., *Polydora* sp., *Sigambra grubii*, and *Melanoides tuberculata* were not detected in the initial impact but were present in the post impact (Table 2). Of the benthic invertebrate taxa sampled in the estuary, most (42%) had significant reductions in their relative abundance or were not sampled (50%) during post impact conditions when compared to the pre-impact assessment. One taxa (*Macoma* sp.) increased in abundance during post-impact conditions (Table 2).

Species diversity (H’) was on average 1.1 ± 0.4 during pre-impact periods, and decreased significantly between years of 2017-2018 (PERMANOVA F= 16.9; p<0.01; Table 2). In January 2020 the diversity levels were similar to the pre impact conditions (Figure 3). The equitability index was only slightly different between pre-impact and post-impact conditions (p=0.04; Table S1), and ranged on average from 0.70 to 0.84 in August 2018 and January 2020, respectively (Figure 3). Species richness (S) decreased markedly from pre-impact to post-impact conditions (PERMANOVA F=14.6, P<0.001), following the 50% reduction in species sampled during the monitoring period (Table 2). In January 2020, the species richness (Margalef index) was similar to pre-impact conditions; when we observed the recolonization of five taxa (*Cirrophorus* sp., Oligochaeta spp., *Telinna* sp., *Tagelus prebeius* and Nemertea; Table 2). Taxa dominance was higher during the low diversity period (2017-2019; range of 0.7 to 0.78) due to the dominance of *Laeonereis* sp., *Macoma* sp., and *Heleobia australis*; and returned to pre-impact levels in January 2020 (Figure 3; Table 2).

**Figure 3.**
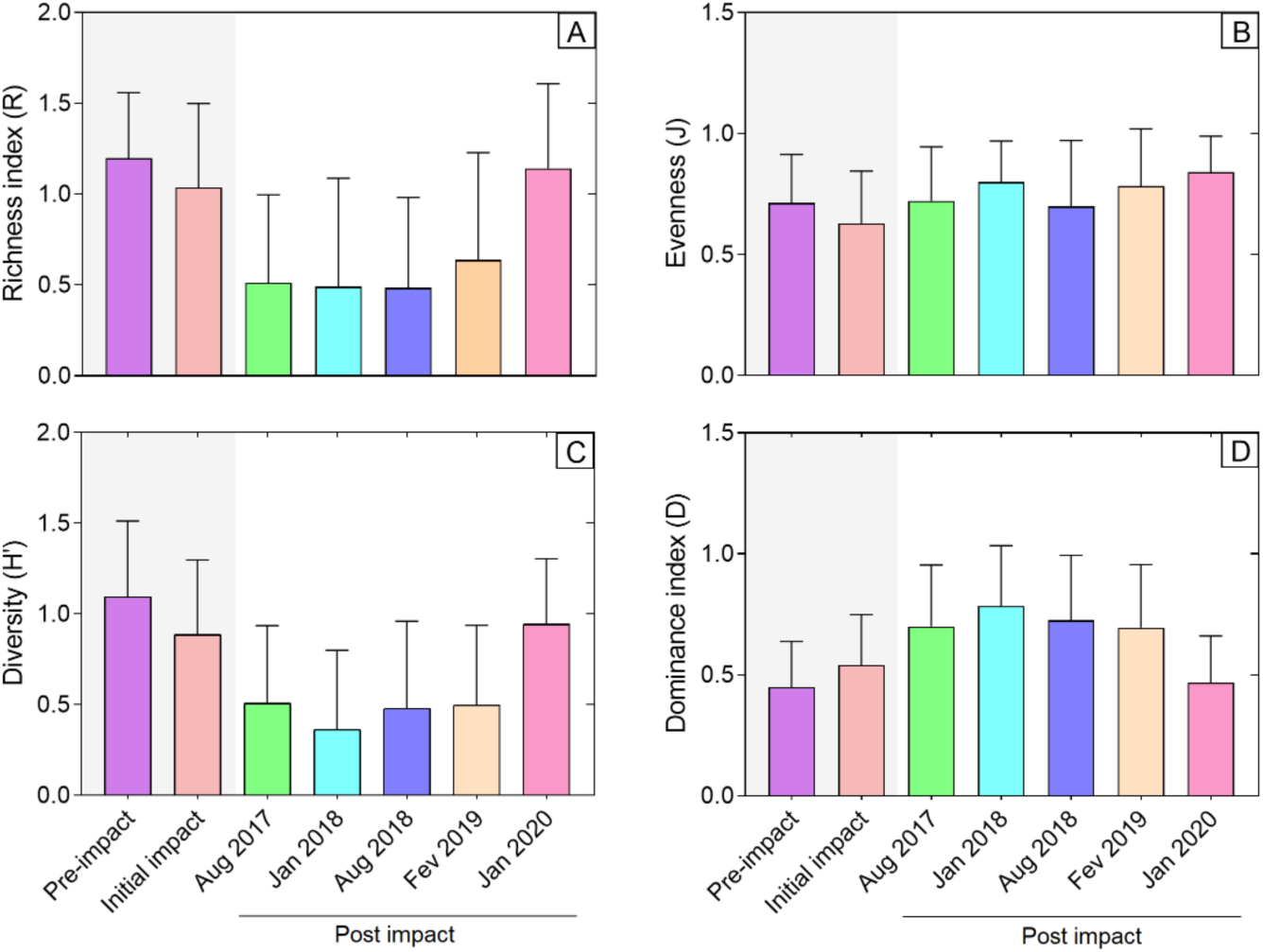
Ecological indicators values in each assessment (pre-impact, initial impact, and post impact). (A) Richness index, (B) Evenness, (C) Shannon-Wiener index, and (D) Dominance index.

### Benthic macrofaunal functional groups

In the Rio Doce estuary, we observed a decrease in surface deposit-feeders (SDF) and an increase in detritivores (DF) after the impact (Figure 2). SDF were dominant in post-impact periods, with a relative abundance of over 70% (Figure 2). During the pre-impact and initial impact, SDF were followed by Omnivorous, Carnivores and Others (OCO; 18.2% and 10.1%, respectively) and detritivores (DF; 10.3% and 8.1%, respectively; Figure 2). During the post impact condition, detritivores were dominant after SDF corresponding to 18.8%; while OCOs represented 8.1%. It is worth mentioning that during the the post impact conditions, there was an absence of suspension-feeders (SF) in 2018 (January and August) and 2019 (February), which returned in January 2020 with an abundance 8 times lower than pre-impact levels.

In general, macrofaunal functional indexes had weak relationship with the impact levels in the Rio Doce estuary. The macrofauna functional richness index (FRic) ranged on average from 2.05 ± 0.67 in the post impact to 2.27 ± 0.78 in the pre-impact, with no difference between both conditions (F = 1.1, p = 0.33; Table S; Fig.4A). In contrast, FEve showed an opposite pattern, with higher values in the post impact (0.39 ± 0.22) when compared to the pre-impact and initial impact conditions (0.29 ± 0.19 and 0.27 ± 0.15, respectively; F = 5.0, p = 0.01; Table S3; Figure 4). In the pre-impact and post impact condition, the mean values of the FDis were similar (0.21 ± 0.14), as well as showing no difference when compared to the initial impact condition (0.16 ± 0.13; F = 1.8, p = 0.17; table S3; Figure 4). On the other hand, FRaoQ showed the lowest value in the pre-impact (0.08 ± 0.05), while the average value for the post impact was 11.08 ± 31.60, reaching in August 2018 the average value of 17.92 ± 45.36 (Fig.4D).

**Figure 4.**
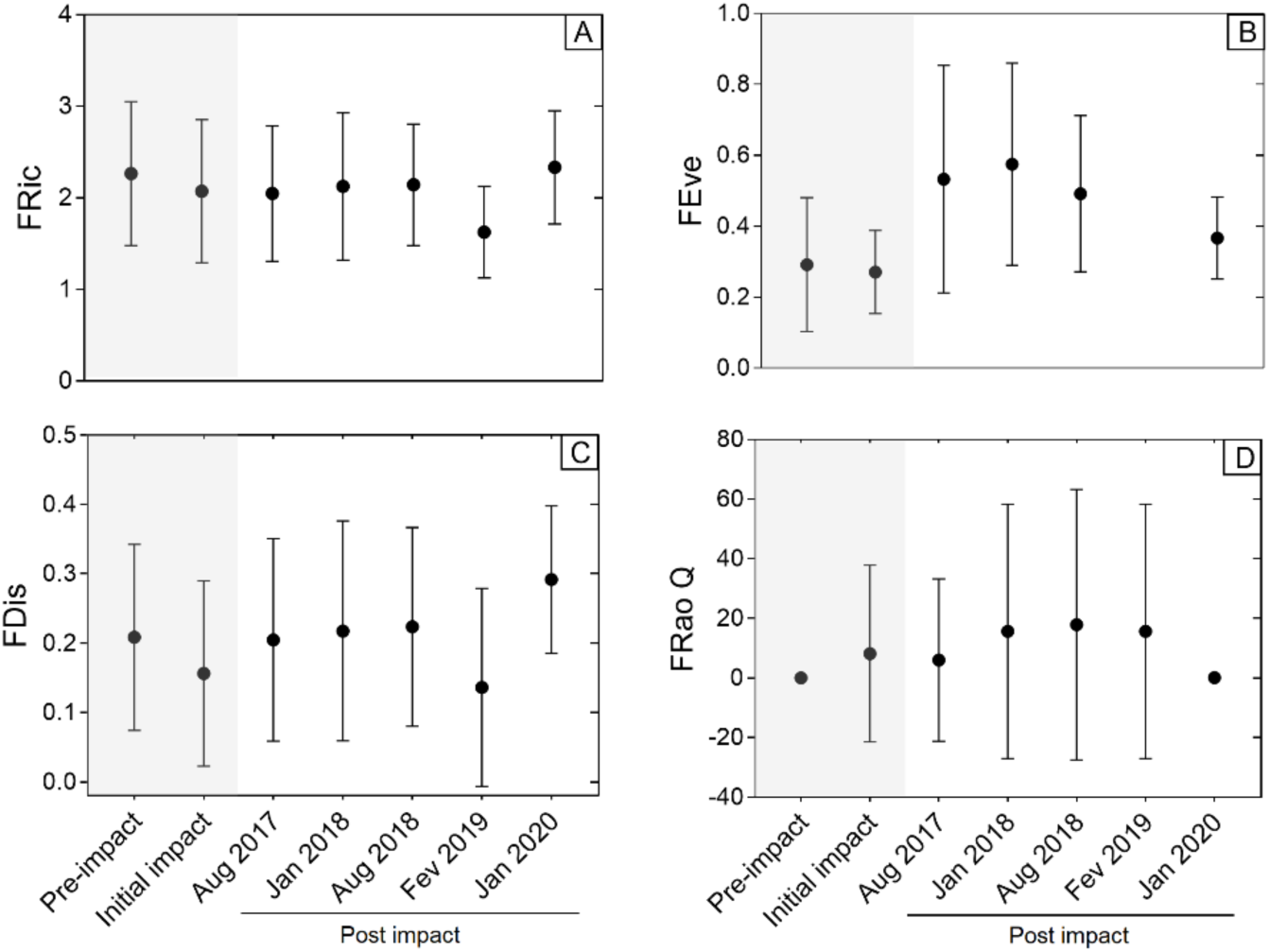
Mean (±SE) macrofaunal functional diversity indices. (A) Functional richness (FRic), (B) functional evenness (FEve), (C) functional dispersion (FDis), and (D) entropy (FRao Q) from assessments on pre-impact, initial impact, and post impact (August 2017, January 2018, August 2018, February 2019, and January 2020) conditions at the Rio Doce estuary.

### Multivariate analyses

The Rio Doce macrofauna was markedly distinct between pre- and post-impact conditions during the 4.2 years of monitoring; which related to the significant decreases in the density organisms present and the number of species not sampled after the impact in 2015 (PERMANOVA F = 40.3; p<0.001; Table S2). The canonical analysis of principal coordinate analysis (CAP) indicated a significant association between the concentrations of potentially toxic elements and the benthic assemblage change (F = 1.99, p = 0.001; Figure 5). The CAP ordination showed that the potentially toxic elements in general are more associated with the post impact condition than the conditions of pre-impact and initial impact. The taxa that mostly contributed to this pattern were *Laeonereis* sp. and *Alitta succinea*, which had a marked density reductions along the study periods and correlated negatively to the high levels of sediment Co, Cu, Ni, and Zn (Co F = 2.45, p = 0.01; Cu F = 1.87, p = 0.05; Ni F = 2.02, p = 0.03; Zn F = 2.58, p = 0.002). Chironomids also had a negative correlation with Cu and positive correlation with Fe and Mn (Fe F = 5.76, p = 0.001; Mn F = 2.05, p = 0.03; Figure 5).

**Figure 5.**
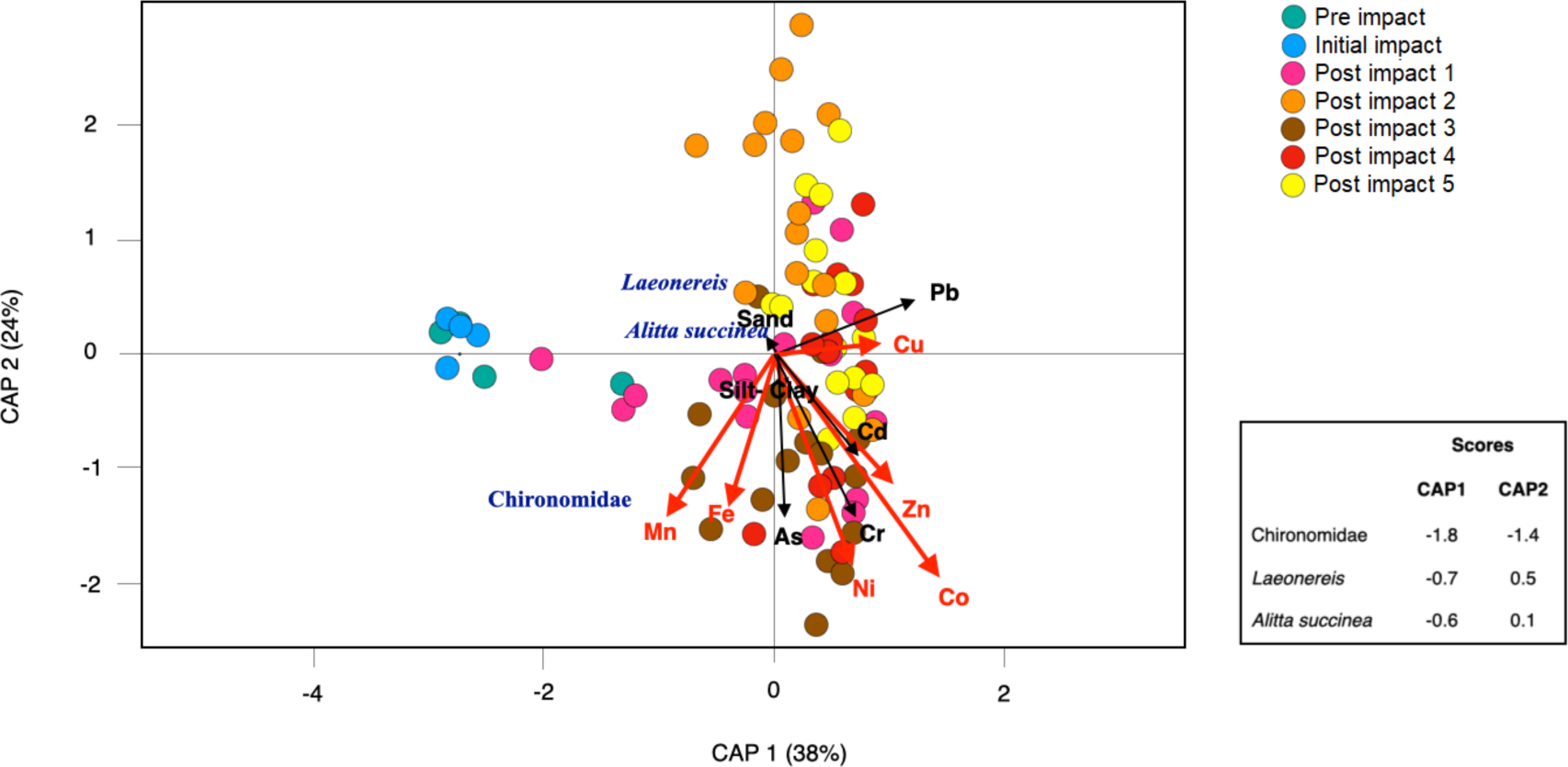
Canonical analyses of principal coordinates (CAP) ordination of samples according to differences in macrofauna density per taxa (number of organisms per m^2^ of sediment) and the contribution of sediment composition (% of sand, and silt+clay) and potentially toxic elements concentration (As, Cd, Co, Cr, Cu, Fe, Mn, Ni, Pb, and Zn) under different assessment conditions (pre-impact, initial impact, and post impact). Vectors are based on Spearman correlation values (p > 0.5) for scores for taxa.

## Discussion

The tailings dam failure and the succeeding arrival of the plume in the Rio Doce estuary increased the sedimentation and accumulation rates, identified by anomalous concentrations of As, Cd, Co, Cr, Cu, Fe, Mn, Ni, Pb, and Zn (Gomes et al., 2017; Queiroz et al., 2021; Barcellos et al., 2022; Queiroz et al., 2022). The concentration of potentially toxic elements in sediments of the Rio Doce estuary reached their highest values during the dry season (i.e., August 2017, August 2018), with concentrations decreasing over time and some seasonal fluctuations. The decrease in contamination levels suggests loss of material through the process of sediment resuspension and transport to the ocean, or the burial of contaminated material (Rudorff et al., 2018; Magris et al., 2019). Additionally, the dissolutive reduction of iron oxyhydroxides in the sediments, the main mineral component of the mine tailings, may also have contributed to this decrease of contamination levels over the years. In the latter case, these data may suggest a release of the studied potentially toxic elements to the estuarine water by bioturbation and natural water irrigation of porewaters (see Barcellos et al., 2022; Queiroz et al., 2021; Queiroz et al., 2022). However, the mean concentrations of As (+21%), Co (+1300%), Cr (+388%), Cu (+377%), Ni (+280%), Pb (+2800%), and Zn (+780%) in January 2020, 4.2 years after the disaster, are still higher than the reference values presented by Gomes et al. (2017). During this period, the sedimentary contamination was frequently and likely constantly above environmental safety thresholds, and caused potentially toxic conditions for benthic macroinvertebrates (Gabriel et al. 2020a). The potentially toxic element concentrations evidence that the Rio Doce estuary has a markedly distinct environmental behavior than freshwater ecosystems in the upper river basin, and that estuarine sediments continue to exhibit a low ecosystem quality even when compared to other polluted urban estuaries in the region (Hadlich et al. 2018).

The Rio Doce estuary sediments are markedly variable spatially but in general are composed of a mixture of fine and sand fractions. These characteristics may favor the accumulation of potentially toxic elements in areas of higher silt-clay fractions, and therefore with variable toxicity (Engel and Fowler, 1979; Heugens et al., 2001). Meanwhile, Cu toxicity s inversely proportional to the total organic matter content (Bryan and Langston, 1992). The low salinity of the Rio Doce estuary is an additional concern related to toxicity, once it influences the partitioning of contaminants between sediments and the overlying waters (Riba et al., 2004). This release has been demonstrated in the Rio Doce estuarine sediments and metals release to porewater was markedly controlled by the irrigation of sub-surface sediments (Barcellos et al., 2021). As a result, we expect that tolerance to toxicity will influence the benthos through reduced richness, recruitment, diversity and altered assemblage composition (Ward and Hutchings 1996; Watzin and Roscigno, 1997; Malmqvist and Hoffsten, 1999; Burd 2002; Venturini et al., 2004). Additionally, sediment contamination by iron in January 2020 is still 324% higher than pre-impact levels, and although often considered an inert element and one of the main elements in the tailings’ composition, iron content has been shown to impact benthic assemblages (Lancellotti and Stotz, 2004).

The high concentrations of potentially toxic elements in the Rio Doce estuary suggests an inhibition in the recovery of the benthic assemblages to their pre-impact state. Although initial impacts included a significant change in sediment grain size and potential effects of rapid sedimentation that are expected from mining waste spills (Gomes et al., 2017; Josefson et al., 2008), our results showed a similar sedimentary texture after the initial mining tailings deposition. The recovery and gradual species succession to the natural state seems then to rely on the removal of the persisting high levels of potentially toxic elements, as the tailings and other organic compounds carried with the mine slurry can have persistent toxicity for some taxa, particularly larger and slow-growing benthic infauna (Burd et al., 2000; Fleeger et al., 2003; Resende et al., 2023). The persisting toxicity in the Rio Doce estuary is similar to other areas contaminated by mine tailings (Olsgard and Hasle; 1993) with strong reduction of taxa richness, species diversity, and density of the benthic macroinvertebrate community in the post impact condition. However, the Rio Doce estuary showed similar levels of species richness when balanced to the lower macrofaunal abundance (Margalef) after the impact, and similar diversity levels at 4.2 years. This suggests that these indexes may mislead the recovery of the benthic fauna since we observed a significant reduction in the number of organisms and many taxa were still absent from the estuary after 4.2 years.

The greatest reduction in diversity and density was seen for annelids. Annelids are classified as the most sensitive fauna to PTE contamination, followed by crustaceans and mollusks (McLusky et al., 1986). *Heleobia australis* and *Laeonereis* sp. were the taxa with the highest densities in the post impact condition. Studies show that the mollusk *H. australis* has tolerance to PTE contamination, tolerates hypoxic/anoxic background conditions, and suggests an opportunistic behavior (Fenchel, 1975; Rakocinski et al., 2000; Echeverría et al., 2010). Likewise, previous studies indicate that the taxa *Laeonereis* sp. shows a bioindicator potential for pollution and stands out due its rapid recolonization in impacted estuaries (Netto and Lana 1994; Omena and Amaral, 2000; Weis et al., 2017). Therefore, our results support the tolerant characteristic of the species that appeared in all impact conditions, although the densities in the post impact condition were lower (Figure 4). A 28-day laboratory study with the Rio Doce sediments compared cores with and without the presence of *Laeonereis* sp. bioturbation and the impact on oxygen and PTE fluxes (Barcellos et al., 2021). One of the major findings was that the benthic bioturbation promoted the oxidation of Fe phases from the mine tailings and prevented the release of PTE bound to the sediments to the aquatic system. The absence of worms in the sediments represented an efflux of Cr, Ni, Cd, and Ba to the aqueous phase. Consequently, the loss of the bioturbation by the benthic community due to deposition of mine tailings at the estuary, may cause an increase in PTE contamination. The *Macoma* sp. (molusk) and *Diastylis* sp. (microcrustacean) taxa showed a different behavior over the sampling period, with an increase in density from the initial impact to the post impact condition, presenting (in certain campaigns) higher densities than those found in the pre-impact. These results may indicate that this taxon is tolerant to PTE contamination. Meanwhile, *Diastylis* sp. was not found since August 2018, suggesting a high sensitivity to PTE contamination, even though this genus is described as early substrate colonizers, highly mobile, which are quickly replaced by more efficient species (Pearson and Rosenberg, 1978; Trannum et al., 2011).

Polychaetes can quickly recolonize tailings and have dominated the estuarine environment 4.2 years post-impact. *Glycera* sp., *Magelona* sp., *Polydora* sp., *Sigambra grubii*, and *Melanoides tuberculata* were found in the post impact condition, after the initial impact, although the detections were punctual and of low abundance, and consequently, low density. These taxa that appeared only in the post impact condition can be considered as an occasional or opportunistic species. The complete absence or low density of opportunistic species in post impact conditions such as capitellids, spionids, and cirratullids polychaetes was notable. This may be due to the effects of the water column because of the tailings resuspension, uninterrupted sedimentation at the estuary bottom, and/or the compact nature of the tailings at the bottom, which may exceed the tolerance levels of these groups (Lancellotti and Stotz, 2004).

The dominance of trophic groups in the Rio Doce estuary in all impacted conditions was structured mainly by surface deposit feeders (SDF) in a similar pattern as observed in other regions impacted by mining tailings (Lancellotti and Stotz, 2004). The predominance of SDF in the impact conditions may be explained by its tolerance to pollution and its high density may prevent the establishment of other organisms from other functional groups (Nunes et al., 2008; Levinton, 1972). In addition, the absence of SF in the post impact condition may be associated with sediment, since the coexistence of SDF and SF depends on a substrate stability that allows suspension feeders to colonize it (Young and Rhoads, 1971). Therefore, the return of suspension feeders (SF) individuals and the decrease in SDF in January 2020, and the variation between omnivores, carnivores and others (OCO), and detritivore feeders (DF) post impact, suggest a slow recovery of the benthic assemblages after 4.2 years of the disaster with seasonal influences (Figure 2B). Regarding the functional metrics, the functional richness (FRic) decreased in the post impact condition (Figure 4), but not as much as the diversity (H’; Figure 3), suggesting that many functions were still present in the Rio Doce estuary even with few species. The post impact condition showed higher FEve values but with less diversity and richness (Figure 3; Figure 4). Although the ecological metrics in the post impact condition are low, their functional characteristics were relatively uniform.

This study has important management implications for the Rio Doce basin. First, it is clear that estuarine ecosystems can become hotspots of sediment contamination after disasters and therefore we can expect decades for the full recovery in the estuarine biodiversity and food chain as previously suggested for the Rio Doce and observed elsewhere (Burd et al., 2000; Burd 2002; Gabriel et al., 2020b). It is also clear that the recovery of water quality in the Rio Doce basin, or in the estuary, is not associated with a decrease in the contamination levels of sedimentary ecosystems and the return of the benthic macrofauna composition to its pre-impact condition. In the same way, our work showed that some, but not all of the taxa previously sampled in the estuary recolonized the impacted areas, suggesting variable levels of tolerance to pollution. This suggests that the use of ecotoxicological essays that demonstrate inert risks of freshwater soils to the selected (model or native) aquatic biota are not applicable estuarine ecosystems impacted by this tragedy and many tests suffer from experimental artifacts (Martinez et al., 2022). Even if levels of toxicity meet accepted guidelines in the Rio Doce estuary, the past, current and future damages to ecosystem services (e.g. fishing, tourism, cultural) to the indigenous communities will never be properly quantified. For the biota, given the high spatial and seasonal variability in the Rio Doce benthos, study designs that can better incorporate both scales will be better positioned to monitor the recovery of this ecosystem; which is likely to occur when all sources of contaminants are removed.

## Acknowledgements

The authors wish to thank all their colleagues involved in the SOLOS-BENTOS Network.

## Declarations

### Credit statement

Angelo Bernardino and Tiago Ferreira conceived the ideas, obtained funding and designed methodology; Gabriel Coppo, Fabricio Gabriel, Hermano Queiroz, Diego Barcellos, Ana Mazzuco and Angelo Bernardino collected the data; Gabriel Coppo, Fabricio Gabriel and Ana Carolina Mazzuco analysed the data; Gabriel Coppo and Angelo Bernardino led the writing of the manuscript. All authors contributed critically to the drafts and gave final approval for publication.

## Supplementary material

**Table S1.** PERMANOVA results of the analysis comparing variations in ecological metrics (macrofauna density, richness, diversity, and evenness) between pre-impact, initial impact and post impact conditions and events in the Rio Doce estuary. Df = Degrees of Freedom; SS = Sum of Squares; MS = Mean of Squares. Significative values (p<0.05) are presented in bold. Note: Insecta was removed and only marine taxa were considered in the analysis.

| Density |  |  |  |  |  |
| --- | --- | --- | --- | --- | --- |
|  | df | SS | MS | F | *p* |
| Impact | 2 | 419.2 | 209.5 | 62.5 | **<0.001** |
| Event | 4 | 40.4 | 10.1 | 3.0 | **0.020** |
| I x E | 4 | 14.2 | 3.6 | 1.1 | 0.378 |
| Residuals | 141 | 472.7 | 3.3 |  |  |
| Richness |  |  |  |  |  |
|  | df | SS | MS | F | *p* |
| Impact | 2 | 7.2 | 3.6 | 14.6 | **<0.001** |
| Event | 4 | 3.7 | 0.9 | 2.8 | **0.006** |
| I x E | 4 | 2.1 | 0.5 | 2.1 | 0.083 |
| Residuals | 141 | 34.5 | 0.2 |  |  |
| Diversity |  |  |  |  |  |
|  | df | SS | MS | F | *p* |
| Impact | 2 | 7.0 | 3.5 | 16.9 | **<0.001** |
| Event | 4 | 3.7 | 0.9 | 4.5 | **0.002** |
| I x E | 4 | 1.6 | 0.4 | 1.9 | **0.010** |
| Residuals | 141 | 29.0 | 0.2 |  |  |
| Evenness |  |  |  |  |  |
|  | df | SS | MS | F | *p* |
| Impact | 2 | 0.6 | 0.3 | 3.3 | **0.042** |
| Event | 4 | 0.4 | 0.1 | 1.0 | 0.402 |
| I x E | 4 | 0.1 | 0.02 | 0.3 | 0.895 |
| Residuals | 141 | 11.1 | 0.1 |  |  |

**Table S2.**
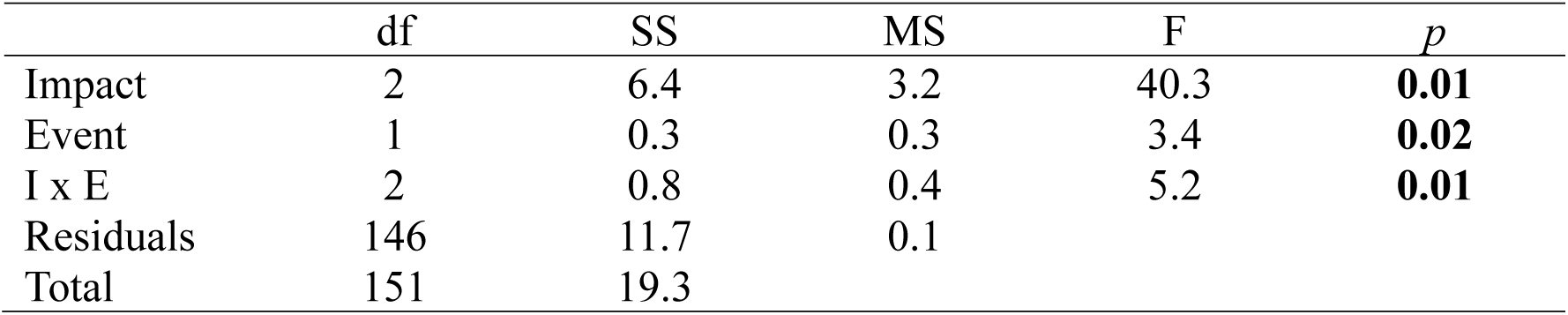
PERMANOVA results comparing variations in macrofaunal assemblages between pre-impact, initial impact and post impact conditions and events in the Rio Doce estuary. Df = Degrees of Freedom; SS = Sum of Squares; MS = Mean of Squares. Significative values (p<0.05) are presented in bold. Note: Insecta was removed and only marine taxa were considered in the analysis.

|  | df | SS | MS | F | *p* |
| --- | --- | --- | --- | --- | --- |
| Impact | 2 | 6.4 | 3.2 | 40.3 | **0.01** |
| Event | 1 | 0.3 | 0.3 | 3.4 | **0.02** |
| I x E | 2 | 0.8 | 0.4 | 5.2 | **0.01** |
| Residuals | 146 | 11.7 | 0.1 |  |  |
| Total | 151 | 19.3 |  |  |  |

**Table S3.** PERMANOVA results of the analysis comparing macrofaunal assemblage functioning (richness FRic, dispersion FDis, evenness FEve, and entropy FRaoQ) between pre-impact, initial impact and post impact conditions and events in the Rio Doce estuary. Df = Degrees of Freedom; SS = Sum of Squares; MS = Mean of Squares. Significative values (p<0.05) are presented in bold. Note: Insecta was removed and only marine taxa were considered in the analysis.

| FRic |  |  |  |  |  |
| --- | --- | --- | --- | --- | --- |
|  | df | SS | MS | F | *p* |
| Impact | 2 | 1.1 | 0.5 | 1.1 | 0.325 |
| Event | 4 | 7.9 | 1.9 | 4.2 | **0.003** |
| I x E | 4 | 5.4 | 1.3 | 2.9 | **0.026** |
| Residuals | 128 | 60.3 | 0.5 |  |  |
| FEve |  |  |  |  |  |
|  | Df | SS | MS | F | *p* |
| Impact | 2 | 0.5 | 0.2 | 5.0 | **0.013** |
| Event | 3 | 0.2 | 0.1 | 1.2 | 0.323 |
| I x E | 3 | 0.01 | 0.003 | 0.1 | 0.970 |
| Residuals | 34 | 1.6 | 0.05 |  |  |
| FDis |  |  |  |  |  |
|  | df | SS | MS | F | *p* |
| Impact | 2 | 0.1 | 0.03 | 1.7 | 0.179 |
| Event | 4 | 0.2 | 0.05 | 2.7 | **0.032** |
| I x E | 4 | 0.04 | 0.01 | 0.5 | 0.715 |
| Residuals | 128 | 2.4 | 0.02 |  |  |
| FRaoQ |  |  |  |  |  |
|  | df | SS | MS | F | *p* |
| Impact | 2 | 0.02 | 0.01 | 1.8 | 0.167 |
| Event | 4 | 0.1 | 0.02 | 3.8 | **0.006** |
| I x E | 4 | 0.01 | 0.003 | 0.5 | 0.748 |
| Residuals | 119 | 0.7 | 0.006 |  |  |

## References

Abessa, D., Burton, G.A., Cervi, E.C., Simpson, S.L., Stubblefield, W., Ribeiro, C.C., Cruz, A.C.F., Kruger, G., Smith, R. (2023). Has the Rio Doce “time bomb” been defused? Using a weight-of-evidence approach to determine sediment quality. Integrated Environmental Assessment and Management, in press.

Aires, U.R.V., Santos, B.S.M., Coelho, C.D., Silva, D.D., & Calijuri, M.L. (2018). Changes in land use and land cover as a result of the failure of a tailings dam in Mariana, MG, Brazil. Land Use Policy, 70:63–70. 10.1016/j.landusepol.2017.10.026

Almeida, C.A., Oliveira, A.F., Pacheco, A.A., Lopes, R.P., Neves, A.A., & Queiroz, M.E.L.R. (2018). Characterization and evaluation of sorption potential of the iron mine waste after Samarco dam disaster in Doce River basin – Brazil. Chemosphere, 209:411–420. 10.1016/j.chemosphere.2018.06.071

Amaral, A.C.Z., & Nonato, E.F. (1996). Annelida Polychaeta: Características, glossário e chaves para famílias e gêneros da costa brasileira. Editora UNICAMP, Campinas, SP.

Amaral, A.C.Z., Rizzo, A.E., & Arruda, E.P. (2005). Manual de identificação dos invertebrados marinhos da região sul-sudeste do Brasil: v.1. Editora da Universidade de São Paulo, São Paulo.

Anderson, M.J. (2017). Permutational multivariate analysis of variance (PERMANOVA) In: Wiley StatsRef: statistics reference online, ©2014–2017. John Wiley & Sons, Ltd. Doi: 10.1002/9781118445112.stat07841.

Anderson, M.J., & Willis, T.J. (2003). Canonical analysis of principal coordinates: a useful method of constrained ordination for ecology. Ecology, 84:511–525. Doi: 10.1890/0012-9658(2003)084[0511:CAOPCA]2.0.CO;2.

Andrades, R., Guabiroba, H.C., Hora, M.S.C., Martins, R.F., Rodrigues, V.L.A., Vilar, C.C., Giarrizzo, T., & Joyeux, J.-C. (2020). Early evidences of niche shifts in estuarine fishes following one of the world’s largest mining dam disasters. Marine Pollution Bulletin 154:111073. 10.1016/j.marpolbul.2020.111073

Andrades, R., Martins, R.F., Guabiroba, H.C., Rodrigues, V.L.A., Szablak, F.T., Bastos, K.V., Bastos, P.G.P., Lima, L.R.S., Vilar, C.C., & Joyeux, J.-C. (2021). Effects of seasonal contaminant remobilization on the community trophic dynamics in a Brazilian tropical estuary. Science of The Total Environment 801:149670. 10.1016/j.scitotenv.2021.149670

Barcellos, D., Jensen, S. S., Bernardino, A. F., Gabriel, F. A., Ferreira, T. O., & Quintana, C. O. (2021). Benthic bioturbation: a canary in the mine for the retention and release of metals from estuarine sediments. Marine Pollution Bulletin 172, 112912.

Barcellos, D., Queiroz, H. M., Ferreira, A. D., Bernardino, A. F., Nóbrega, G. N., Otero, X. L., & Ferreira, T. O. (2022). Short-term Fe reduction and metal dynamics in estuarine soils impacted by Fe-rich mine tailings. Applied Geochemistry 136:105134.

Bernardino, A.F., Netto, S.A., Pagliosa, P.R., Barros, F., Christofoletti, R.A., Rosa-Filho, J.S., Colling, J., & Lana, P.C. (2015). Predicting ecological changes on benthic estuarine assemblages through decadal climate trends along Brazilian marine ecoregions. *Estuarine*, Coastal and Shelf Science 166:74–82. 10.1016/j.ecss.2015.05.021

Bernardino, A.F., Pais, F.S., Oliveira, L.S., Gabriel, F.A., Ferreira, T.O., Queiroz, H.M., & Mazzuco, A.C. (2019). Chronic trace metals effects of mine tailings on estuarine assemblages revealed by environmental DNA. PeerJ 7(7):e8042. 10.7717/peerj.8042

Botta-Dukát, Z. (2005). Rao’s quadratic entropy as a measure of functional diversity based on multiple traits. Journal of vegetation Science 16:533–540. Doi: 10.1111/j.1654-1103.2005.tb02393.x.

Bryan, G.W., & Langston, W.J. (1992). Bioavailability, accumulation and effects of heavy metals in sediments with a special reference to United Kingdom estuaries: a review. Environmental Pollution 76:89–131

Bryant, V., Newbery, D.M., McLusky, D.S., & Campbell, R. (1985). Effect of temperature and salinity on the toxicity of nickel and zinc to two estuarine invertebrates (*Corophium volutator*, Macoma balthica). Marine Ecology Progress Series 24:139–153. http://www.jstor.org/stable/24816860

Burd, B. (2002). Evaluation of mine tailings effects on a benthic marine infaunal community over 29 years. Marine Environmental Research 53(5):481–519. 10.1016/S0141-1136(02)00092-2

Burd, B., Macdonald, R., & Boyd, J. (2000). Punctuated recovery of sediments and benthic infauna: a 19 year study of tailings deposition in a British Columbia fjord. Marine Environmental Research 49(2):145–175. 10.1016/S0141-1136(99)00058-6

Clarke, K.R., & Gorley, R.N. (2006). PRIMER v6: User Manual/Tutorial. 6 ed. Plymouth.

Clements, W.H., Carlisle, D.M., Lazorchak, J.M., & Johnson, P.C. (2000). Heavy metals structure benthic communities in Colorado mountain streams. Ecological Applications 10:626–638. 10.1890/1051-0761(2000)010[0626:HMSBCI]2.0.CO;2

Coppo, G., Pais, F.S., Ferreira, T.O., Halanych, K.M., Donnelly, K., Mazzuco, A.C., & Bernardino, A.F. (2023). Transition of an estuarine benthic meiofauna assemblage 1.7 and 2.8 years after a mining disaster. PeerJ 11:e14992. 10.7717/peerj.14992

Dean, W.E. (1974). Determination of carbonate and organic matter in calcareous sediments and sediments rocks by loss on ignition: comparison with other methods. SEPM Journal of Sedimentary Research. DOI: 44:242–248. 10.1306/74d729d2-2b21-11d7-8648000102c1865d

Do Carmo, F.F., Kamino, L.H.Y., Tobias Junior, R., Campos, I.C., Carmo, F.F., Silvino, G., Castro, K.J.S.X., Mauro, M.L., Rodrigues, N.U.A., Miranda, M.P.S., & Pinto, C.E.F. (2017). Fundão tailings dam failures: the environment tragedy of the largest technological disaster of Brazilian mining in global context. Perspectives in Ecology and Conservation 15(3):145–151. 10.1016/j.pecon.2017.06.002

Echeverría, C.A., Neves, R.A.F., Pessoa, L.A., & Paiva, P.C. (2010). Spatial and temporal distribution of the gastropod Heleobia australis in an eutrophic estuarine system suggests a metapopulation dynamics. Natural Science 2(8):860–867. 10.4236/ns.2010.28108

Engel, D.W., & Fowler, B.A. (1979). Factors influencing cadmium accumulation and its toxicity to marine organisms. Environmental Health Perspectives 28:81–88

Fenchel, T. (1975). Factors determining the distribution patterns of mud snails (Hydrobiidae). Oecologia 20(1), 1–17.

Fisher, R.A., Corbet, A.S., & Williams, C.B. (1943). The relation between the number of species and the number of individuals in a random sample of animal population. Journal of Animal Ecology 12:42–58.

Fleeger, J.W., Carman, K.R., & Nisbet, R.M. (2003). Indirect effects of contaminants in aquatic ecosystems. Science of The Total Environment 317(1-3):207–233. 10.1016/S0048-9697(03)00141-4

Gabriel, F.A., Ferreira, A.D., Queiroz, H.M., Vasconcelos, A.L.S., Ferreira, T.O., & Bernardino, A.F. (2021). Long-term contamination of the Rio Doce estuary as a result of Brazil’s largest environmental disaster. Perspectives in Ecology and Conservation 19(4):417–428. 10.1016/j.pecon.2021.09.001

Gabriel, F.A., Hauser-Davis, R.A., Soares, L., Mazzuco, A.C.A., Rocha, R.C.C., Saint Pierre, T.D., Saggioro, E., Correia, F.V., Ferreira, T.O., & Bernardino, A.F. (2020b). Contamination and oxidative stress biomarkers in estuarine fish following a mine tailing disaster. PeerJ 8:e10266. 10.7717/peerj.10266

Gabriel, F.A., Silva, A.G., Queiroz, H.M., Ferreira, T.O., Hauser-Davis, R.A., & Bernardino, A.F. (2020a). Ecological risks of metal and metalloid contamination in the Rio Doce estuary. Integrated Environmental Assessment and Management16(5):655–660. 10.1002/ieam.4250

Gee, G.W., & Bauder, J.W. (1986). Particle-size analysis. In: A. Klute (Ed.), Methods soil Anal Part 1 - Physical Mineral Methods (2nd ed., pp.383–411). American Society of Agronomy.

Gnandi, K., Tchangbedji, G., Killi, K., Baba, G., & Abbe, K. (2006). The Impact of Phosphate Mine Tailings on the Bioaccumulation of Heavy Metals in Marine Fish and Crustaceans from the Coastal Zone of Togo. Mine Water Environment 25:56–62. 10.1007/s10230-006-0108-4

Gomes, L.E.O., Correa, L.B., Sa, F., Neto, R.R., & Bernardino, A.F. (2017). The impacts of the Samarco mine tailing spill on the Rio Doce estuary, Eastern Brazil. Marine Pollution Bulletin 120(1–2):28–36. 10.1016/j.marpolbul.2017.04.056

Hadlich, H.L., Venturini, N., Martins, C.C., Hatje, V., Tinelli, P., Gomes, L.E.O., & Bernardino, A.F. (2018). Multiple biogeochemical indicators of environmental quality in tropical estuaries reveal contrasting conservation opportunities. Ecological Indicators 95(1): 21–31. 10.1016/j.ecolind.2018.07.027

Heugens, E.H.W., Hendriks, A.J., Dekker, T., van Straalen, N.M., & Admiraal, W. (2001). A review of the effects of multiple stressors on aquatic organisms and analysis of uncertainty factors for use in risk assessment. Critical Reviews in Toxicology 31(3):247–284. 10.1080/20014091111695

Hurlbert, S.H. (1971). The nonconcept of species diversity: a critique and alternative parameters. Ecology 52: 577–586.

Hyland, J., Balthis, L., Karakassi, I., Magni, P., Petrov, A., Shine, J., Vestergaard, O., & Warwick, R. (2005). Organic carbon content of sediments as an indicator of stress in the marine benthos. Marine Ecology Progress Series 295:91–103. 10.3354/meps295091

Jaishankar, M., Tseten, T., Anbalagan, N., Matthew, B.B., & Beeregowda, K.N. (2014). Toxicity, mechanism and health effects of some heavy metals. Interdisciplinary Toxicology 7(2):60–72. 10.2478/intox-2014-0009

Josefson, A.B., Hansen, J.L.S., Asmund, G., & Johansen, P. (2008). Threshold response of benthic macrofauna integrity to metal contamination in West Greenland. Marine Pollution Bulletin 56(7):1265–1274. 10.1016/j.marpolbul.2008.04.028

Kraus, U., & Wiegand, J. (2006). Long-term effects of the Aznalcóllar mine spill - heavy metals content and mobility in soils ans sediments of the Guadiamar river valley (SW Spain). Science of The Total Environment 367(2-3):855–871. 10.1016/j.scitotenv.2005.12.027

Kristensen, E., Delefosse, M., Quintana, C.O., Flindt, M.R., & Valdemarsen, T. (2014). Influence of benthic macrofauna community shifts on ecosystem functioning in shallow estuaries. Frontiers in Marine Science 1(41). 10.3389/fmars.2014.00041

Kristensen, E., Penha-Lopes, G., Delefosse, M., Valdemarsen, T., Quintana, C.O., & Banta, G.T. (2012). What is bioturbation? Need for a precise definition for fauna in aquatic science. Marine Ecology Progress Series 446: 285. 10.3354/meps09506

Lahijanzadeh, A.R., Rouzbahani, M.M., Sabzalipour, S., & Nabavi, S.M.B. (2019). Ecological risk of potentially toxic elements (PTEs) in sediments, seawater, wastewater, and benthic macroinvertebrates, Persian Gulf. Marine Pollution Bulletin 145:377–389. 10.1016/j.marpolbul.2019.05.030

Lancellotti, D.A., & Stotz, W.B. (2004). Effects of shoreline discharge of iron mine tailings on a marine soft-bottom community in northern Chile. Marine Pollution Bulletin 48:303–312. 10.1016/j.marpolbul.2003.08.005

Levinton, J. (1972). Stability and trophic structure in deposit-feeding and suspension-feeding communities. The American Naturalist 106(950):472–486

Magris, R.A., Marta-Almeida, M., Monteiro, J.A.F., & Ban, N.C. (2019). A modelling approach to assess the impact of land mining on marine biodiversity: assessment in coastal catchments experiencing catastrophic events (SW Brazil). Science of The Total Environment 659: 828–840. 10.1016/j.scitotenv.2018.12.238

Malmqvist, B., & Hoffsten, P.-O. (1999). Influence of drainage from old mine deposits on benthic macroinvertebrate communities in central Swedish streams. Water Research 33(10):2415–2423.

Margalef, R. (1958). Information Theory in Ecology. General Systems 3:36–71.

Mason, N.W.H., Mouillot, D., Lee, W.G., & Wilson, J.B. 2005. Functional richness, functional evenness and functional divergence: the primary components of functional diversity. Oikos 111(1):112–118. Doi: 10.1111/j.0030-1299.2005.13886.x.

Martinez, A.S., Underwood, T., Christofoletti, R.A., Pardal, A., Fortuna, M.A., Marelo-Silva, J., Morais, G.C., Lana, P.C. (2022). Reviewing the effects of contamination on the biota of Brazilian coastal ecosystems: scientific challenges for a developing country in a changing world. Science of the Total Environment, 803: 150097

McLusky, D.S., Bryant, V., & Campbell, R. (1986). The effects of temperature and salinity on the toxicity of heavy metals to marine and estuarine invertebrates. Oceanography and Marine Biology: An Annual Review 24:481–520.

Muniz, P., Venturini, N., Hutton, M., Kandratavicius, N., Pita, A., Brugnoli, E., Burone, L., & García-Rodríguez. F. (2011). Ecosystem health of Montevideo coastal zone: A multi approach using some different benthic indicators to improve a ten-year-ago assessment. Journal of Sea Research 65(1): 38–50. 10.1016/j.seares.2010.07.001

Muniz, P., Venturini, N., Martínez, A. (2002). Physico-chemical and pollutants of the benthic environment of the Montevideo Coastal Zone, Uruguay. Marine Pollution Bulletin 44:962–968.

Netto, S.A., & Lana, P.C. (1994). Effects of sediment disturbance on the structure of benthic fauna in a subtropical tidal creek of southeastern Brazil. Marine Ecology Progress Series 106:239–247

Nunes, M., Coelho, J.P., Cardoso, P.G., Pereira, M.E., Duarte, A.C., & Pardal, M.A. (2008). The macrobenthic community along a mercury contamination in a temperate estuarine system (Ria de Aveiro, Portugal). Science of The Total Environment 405(1-3):186–94. 10.1016/j.scitotenv.2008.07.009

Okindele, E.O., Omisakin, O.D., Oni, O.A., Aliu, O.O., Omoniyi, G.E., & Akinpelu, O.T. (2020). Heavy metal toxicity in the water column and benthic sediments of a degraded tropical stream. Ecotoxicology and Environmental Safety 190:110153. 10.1016/j.ecoenv.2019.110153

Olsgard, F., & Hasle, J.R. (1993). Impact of waste from titanium mining on benthic fauna. Journal of Experimental Marine Biology and Ecology 172(1-2):185–213. 10.1016/0022-0981(93)90097-8

Omena, E.P., & Amaral, A.C.Z. (2000). Population dynamics and secondary production of Laeonereis acuta (Treadwell, 1923) (Nereididae: Polychaeta). Bulletin of Marine Science 67:421-431

Pearson, T.H., & Rosenberg, R. (1978) Macrobenthic Succession in Relation to Organic Enrichment and Pollution of the Marine Environment. Oceanography and Marine Biology: An Annual Review 16:229–311.

Queiroz, H. M., Ying, S. C., Bernardino, A. F., Barcellos, D., Nóbrega, G. N., Otero, X. L., &. Ferreira, T. O. (2021a). Role of Fe dynamic in release of metals at Rio Doce estuary: unfolding a mining disaster. Marine Pollution Bulletin 166:112267.

Queiroz, H. M., Ying, S. C., Abernathy, M., Barcellos, D., Gabriel, F. A., Otero, X. L., … & Ferreira, T. O. (2021b). Manganese: The overlooked contaminant in the world largest mine tailings dam collapse. Environment International 146: 106284.

Queiroz, H. M., Ferreira, T. O., Barcellos, D., Nóbrega, G. N., Antelo, J., Otero, X. L., & Bernardino, A. F. (2021c). From sinks to sources: The role of Fe oxyhydroxide transformations on phosphorus dynamics in estuarine soils. Journal of Environmental Management 278:111575.

Queiroz, H. M., Ruiz, F., Deng, Y., de Souza Júnior, V. S., Ferreira, A. D., Otero, X. L., … & Ferreira, T. O. (2022). Mine tailings in a redox-active environment: Iron geochemistry and potential environmental consequences. Science of The Total Environment 807: 151050.

Queiroz, H.M., Nóbrega, G.N., Ferreira, T.O., Almeida, L.S., Romero, T.B., Santaella, S.T., Bernardino, A.F., & Otero, X.L. (2018). The Samarco mine tailing disaster: a possible time-bomb for heavy metals contamination? Science of The Total Environment 637–638:498-506. 10.1016/j.scitotenv.2018.04.370

R Core Team. (2021). R: A Language and Environment for Statistical Computing. R Foundation for Statistical Computing, Vienna, Austria. http://www.R-project.org/.

Rakocinski, C.F., Brown, S.S., Gaston, G.R., Heard, R.W., Walker, W.W. & Summers, J.K. (2000). Species-abun-dance-biomass responses by estuarine macrobenthos to sediment chemical contamination. Journal of Aquatic Ecosystem Stress and Recovery 7(3):201–214.

Resende, J.S.S., Pereira, R., Bernardino, A.F., Longhini, C.M., Lehrback, B.D., Silva, C.A., Costa, E.S., Elvert, M., Neto, R.R. (2023). Organic matter changes at the Rio Doce river mouth caused by the Fundao Dam mine tailing collapse. Water Air Soil Pollution, 234: 486

Reynoldson, T.B. (1987). Interactions between sediment contaminants and benthic organisms. Hydrobiologia 149:53–66.

Riba, I., Del Valls, T.Á., Forja, J.M., & Gómez-Parra, A. (2004). The influence of pH and salinity on the toxicity of heavy metals in sediment to the estuarine clam Ruditapes philippinarum. Environmental Toxicology and Chemistry 23(5):1100–1107.

Riba, I., DelValls, T.A., Forja, J.M., & Gómez-Parra, A. (2002). Evaluating the Heavy Metal Contamination in Sediments from the Guadalquivir Estuary after the Aznalcóllar Mining Spill (SW Spain): A Multivariate Analysis Approach. Environmental Monitoring and Assessment 77:191–207. 10.1023/A:1015828020313

Richard, E. C., Duarte Jr, H. A., Estrada, G. C. D., Bechtold, J., Maioli, B.G., de Freitas, A. H. A., Warner, K. E., & Figueiredo, L. H. M. (2020). Influence of Fundão Tailings Dam Breach on Water Quality in the Doce River Watershed. Integrated Environmental Assessment and Management, 16(5), 583–595. 10.1002/ieam.4311.

Rudorff, N., Rudorff, C.M., Kampel, M., & Ortiz, G. (2018). Remote sensing monitoring of the impact of a major mining wastewater disaster on the turbidity of the Doce River plume off the eastern Brazilian coast. ISPRS Journal of Photogrammetry and Remote Sensing 145(B), 349–361. 10.1016/j.isprsjprs.2018.02.013

Sá, F., Longhini, C.M., Costa, E.S., Silva, C.A., Cagnin, R.C., Gomes, L.E.O., Lima, A.T., Bernardino, A.F., & Neto, R.R. (2021). Time-sequence development of metal(loid)s following the 2015 dam failure in the Doce river estuary, Brazil. Science of The Total Environment 769:144532. 10.1016/j.scitotenv.2020.144532

Silva, C.A., Zacché, D.S., Lehrback, B.D., Cagnin, R.C., Costa, E.S., Longhini, C.M., Bernardino, A.F., Sá, F., & Neto, R.R. (2023). PAHs in estuarine sediments as a consequence of the mine tailings remobilization and transport in the Rio Doce basin. Integrated Environmental Assessment and Management. 10.1002/ieam.4831

Snelgrove, P.V.R., Austen, M.C., Boucher, G., Heip, C., Hutchings, P.A., King, G.M., Koike, I., Lambshead, P.J.D., & Smith, C.R. (2000). Linking Biodiversity Above and Below the Marine Sediment–Water Interface. BioScience 50:1076–1088. 10.1641/0006-3568(2000)050[1076:LBAABT]2.0.CO;2

Sweetman, A.K., Haugland, B.T., Kvassnes, A.J.S., & Bolam, S.G. (2020). Impeded Macrofaunal Colonization and Recovery Following Marine Deposition of Inert and Organically Modified Mine-Tailings. Frontiers in Marine Sciences 7:649. 10.3389/fmars.2020.00649

Trannum, H.C., Olsgard, F., Skei, J.M., Indrehus, J., Øverås, S., Eriksen, J. (2004). Effects of copper, cadmium and contaminated harbour sediments on recolonisation of soft-bottom communities. Journal of Experimental Marine Biology and Ecology 310(1):87–114. 10.1016/j.jembe.2004.04.003

Underwood, A.J., 1992. Beyond BACI: the detection of environmental impact on populations in the real, but variable, world. Journal of Experimental Marine Biology and Ecology 161(2):145–178. 10.1016/0022-0981(92)90094-Q

United States Environmental Protection Agency – USEPA. (1997). Method 3052: Microwave Assisted Acid Digestion of Siliceous and Organically Based Matrices. EPA SW-846.Washington DC.

Varzim, C.S., Hadlich, H.L., Andrades, R., Mazzuco, A.C.A., & Bernardino, A.F. (2019). Tracing pollution in estuarine benthic organisms and its impacts on food webs of the Vitoria Bay estuary. Estuarine, Coastal and Shelf Science 229:10641. 10.1016/j.ecss.2019.106410

Venturini, N., Muniz, P., & Rodriguez, M. (2004). Macrobenthic subtidal communities in relation to sediment pollution: a test of the applicability of the phylum-level meta-analysis approach in a southeastern coastal region of South America. Marine Biology 144:119–126.

Ward, T.J., & Hutchings, P.A. (1996). Effects of trace metals on infaunal species composition in polluted intertidal and subtidal marine sediments near a lead smelter, Spencer Gulf, South Australia. Marine Ecology Progress Series 135:123–135.

Warwick, R.M., & Clarke, K.R. (1993). Comparing the severity of disturbance: a metaanalysis of marine macrobenthic community data. Marine Ecology Progress Series 92(3):221–231.

Watzin, M.C., & Roscigno, P.R. (1997). The effects of zinc contamination on the recruitment and early survival of benthic invertebrates in an estuary. Marine Pollution Bulletin 34(6):443–455. 10.1016/S0025-326X(96)00132-4

Weber, A.A., Sales, C.F., Faria, F.S., Melo, R.M.C., Bazzoli, N., & Rizzo, E. (2020). Effects of metal contamination on liver in two fish species from a highly impacted neotropical river: a case study of the Fundão dam, Brazil. Ecotoxicology and Environmental Safety 190:110–165. 10.1016/j.ecoenv.2020.110165

Weis, W.A., Soares, C.H.L., Quadros, D.P.C., Scheneider, M., & Pagliosa, P.R. (2017). Urbanization effects on different biological organization levels of an estuarine polychaete tolerant to pollution. Ecological Indicators 73:698–707. 10.1016/j.ecolind.2016.10.029

Young, D.K., & Rhoads, DC. (1971). Animal–sediment relations in Cape Cod Bay Massachusetts. Marine Biology 11:242–254

Zhou, Q., Zhang, J., Fu, J., Shi, J., & Jiang, G. (2008). Biomonitoring: An appealing tool for assessment of metal pollution in the aquatic ecosystem. Analytica Chimica Acta 606(2):135–150. 10.1016/j.aca.2007.11.018

